# Characterization of a core fungal community and captivity-induced “mycobiome” change in Fowler’s Toad (*Anaxyrus fowleri*)

**DOI:** 10.1101/2025.07.08.659677

**Authors:** Alexander J Bradshaw, Sinlan Poo, Tracey E. Malter, Rylie M. Strasbaugh, Brianna Bodner, Madison R. Hincher, Anne Devan-Song, Javier F. Tabima

## Abstract

Amphibious animals, such as frogs, are found at the intersection of aquatic and terrestrial ecosystems. They often serve as keystone and sentinel species, essential in nutrient cycling and food webs. In recent decades, amphibians have experienced drastic population declines due to habitat loss, climate change, and disease. These declines have prompted investments in *ex situ* conservation and captive breeding programs, which aim to reduce extinction risk by creating assurance colonies and reintroducing individuals once threats are mitigated. A critical component of these programs is proper husbandry, which ensures the health and longevity of captive populations and their ability to produce offspring that can be reintroduced into the wild. The artificial environment in captivity can profoundly impact animal behavior and health, particularly in relation to diet and nutrition. Diet not only provides nutrients and energy but also shapes the host’s gut microbial community, which in turn impacts digestive health. Complex microbial communities, collectively known as the microbiome, are characterized by the high biodiversity of prokaryotes, microscopic fungi, and viruses. The diet-associated microbiome is increasingly studied for its role in captive animal health and behavior, although research has focused more on bacteria than fungal communities, or the “mycobiome”. Here, we investigated the core mycobiome using metabarcoding of fungal communities in 15 wild-caught *Anaxyrus fowleri* (Fowler’s Toad), documenting shifts as toads transitioned from wild to captive settings. We identified a core set of fungal taxa and observed distinct changes in non-core fungi associated with dietary differences. These findings highlight the dynamic nature of the amphibian mycobiome and the significant impact captivity can have on microbial composition, providing a framework for understanding the role of the amphibian mycobiome in future conservation efforts.

## Introduction

Amphibians play a crucial role in maintaining ecosystem balance, serving as both predators and prey within aquatic and terrestrial food webs and acting as indicators of ecosystem health (Kiesecker *et al*., 2004). However, global populations of amphibians are experiencing drastic declines (Alroy, 2015; Luedtke *et al*., 2023), with estimates suggesting extinction rates exceed more than 200 times the natural background levels (Blaustein *et al*., 2011) and exceeds fossil record extinction by orders of magnitude (Alroy, 2015; Tietje & Rödel, 2018). These declines result from habitat destruction, climate change, pollution, and the emergence of infectious diseases, such as chytridiomycosis caused by the fungus *Batrachochytrium dendrobatidis* (Alford *et al*., 2001; Lips, 2016). While some amphibian species remain abundant, nearly 40% are now classified as threatened by the International Union for Conservation of Nature (IUCN), making them one of the most at-risk vertebrate groups. Conservation efforts, including habitat restoration (Burrow & Lance, 2022) and captive breeding programs (Karlsdóttir *et al*., 2021), have been increasingly prioritized in response to these declines. As conservation strategies are implemented, increasing attention is being paid to the physiological effects of diet and the changing microbiome on animals in activity (Trevelline *et al*., 2019). However, significant gaps remain in understanding how captivity affects amphibian health and microbiome stability (Kueneman *et al*., 2022; Korpita *et al*., 2023).

Microbial communities play essential roles as symbionts across the trophic structure of the natural environment. Microbial communities within the gastrointestinal (GI) tract can be incredibly diverse, including representation across all domains of life, and interact dynamically with the host (Berg *et al*., 2020). The microbiome has been shown to influence the overall health of a host, including dietary preferences and even neurobiological development (Sharon *et al*., 2016; Trevelline & Kohl, 2022). However, the composition of these microbial communities is not static. The GI-associated microbiome can shift in response to dietary alterations and environmental changes, with potentially drastic consequences for host health and adaptation (Diaz & Reese, 2021).

Fungi represent one of the most diverse and ecologically important groups of organisms on Earth, occupying nearly every conceivable habitat and forming complex symbiotic relationships across the tree of life. This immense biodiversity is matched by a remarkable array of biochemical capabilities, making fungi key players in shaping ecosystem function and host physiology. Fungi are well-known for forming mutualistic and parasitic relationships with animals, with some exerting a profound influence on host behavior. For instance, *Cordyceps*, *Entomophthora,* and *Massospora* infect insect hosts and manipulate their behavior to facilitate fungal reproduction (Boyce *et al*., 2019; Loreto & Hughes, 2019). Fungi are environmental engineers, so it is unlikely that within the amphibian gut environment, fungi are passive residents. Rather, GI-associated microfungi could be active participants in shaping host health, digestion, and potentially even behavior. The amphibian gut mycobiome represents a biologically and chemically rich ecosystem whose role in host biology remains poorly understood and warrants deeper exploration.

Early studies suggest that mycobiomes play pivotal roles in host metabolism, the degradation of complex carbohydrates, pathogen resistance, immune system regulation, and also synergistically aid in the spread of both micro and macro fungi (Wagg *et al*., 2019; Harrison *et al*., 2020; Bradshaw *et al*., 2022; Weinstein *et al*., 2022). However, fungal diversity directly influences these functions, making it crucial to characterize and compare fungal communities across environmental gradients. This knowledge gap is particularly concerning in the case of amphibians, which are known to interact with fungi, producing divergent health outcomes in host populations. For instance, the chytrid fungus *B. dendrobatidis* is tolerated by some amphibians (Woodhams *et al*., 2011; Peterson & McKenzie, 2014), but has led to the severe decline or loss of other amphibian species (Longcore *et al*., 1999; Fisher *et al*., 2009; Van Rooij *et al*., 2015; Li *et al*., 2021).

Captive rearing presents a particularly striking example of environmental disruption, where animals experience significant dietary and habitat modifications. Bringing wild animals into captivity presents several challenges, as the captive environment can be significantly different from what the animals are evolutionarily adapted to and often lacks the cues, resources, and complexity of a natural environment (Mason, 2010). These differences in environment can result in variations in behavior (Kelly *et al*., 2025), morphology (O’Regan & Kitchener, 2005; Zack *et al*., 2022), physiology (Turko *et al*., 2023), emerging infections (Taylor *et al*., 1999), and untold other aspects of captive animals compared to their wild counterparts. While some species can adapt to captive environments, others respond negatively to the loss of these natural cues (Mason, 2010). Moreover, captive effects can be compounded throughout multiple generations (Elsbeth McPhee, 2004). The challenges are reversed when a captive-bred or captive-reared animal is released or returned to the wild. While there are known differences between wild and captive animals, many aspects of this environmental change are yet to be explored (Turko *et al*., 2023). In particular, there is a paucity of information regarding the transition wild animals undergo when they are brought into captivity, and, similarly, the transition that captive animals experience when they are released into natural habitats as part of reintroduction or recovery efforts. One significant difference between captive and wild environments is the complexity and variety of food items available to them. While animals in the wild generally consume a wide range of food types, captive animals are often limited to only a few food choices (Turko *et al*., 2023), with supplemental nutrients added to balance out their diets as needed. The reduction or restriction in diet can affect various aspects of an animal’s development (Mitchell *et al*., 2021), physiology, gut microbiota (McKenzie *et al*., 2017; Dallas & Warne, 2023), and behavior (Mason, 2010; Kelleher *et al*., 2019, 2022). Here, we performed metamplicon sequencing of subunit 1 of the Internal transcribed spacer (ITS1) region. ITS1 was chosen over the second subunit (ITS2) due to its tendency to recover higher levels of diversity in non-dikarya microfungal phyla (Blaalid *et al*., 2013). Sequencing was performed on 53 fecal samples from fifteen Fowler’s toads (*Anaxyrus fowleri*) across four weeks to generate a profile of the total mycobiome found in wild animals and their mycobiome community as they transition from wild to captive environments. Furthermore, we also document the overall alpha and beta diversity of our fecal samples, as well as the ecological guild to which the identified fungal taxa belong. This work aims to establish a baseline understanding of the core mycobiome associated with an amphibian, in addition to documenting the total community shifts that occur when these animals are transitioned from the wild to captivity. Documenting the wild and post-captivity mycobiome present in these animals will provide a translational framework for assessing the overall microbiome health of captive amphibians, informing future conservation efforts for threatened and endangered species.

## Results

### Animal capture numbers

Fifteen Fowler’s toads were hand-captured in May of 2022. Each specimen was sexed, weighed, and measured upon initial collection. Seven females, six males, and two toads that were not sexed were collected. Fifteen fecal samples from wild toads were collected before the toads were brought into captivity. In captivity, fecal pellets were collected weekly from both male and female toads, resulting in a total of 53 samples, which were used for downstream DNA extraction and sequencing (Figure 1).

**Figure 1.**
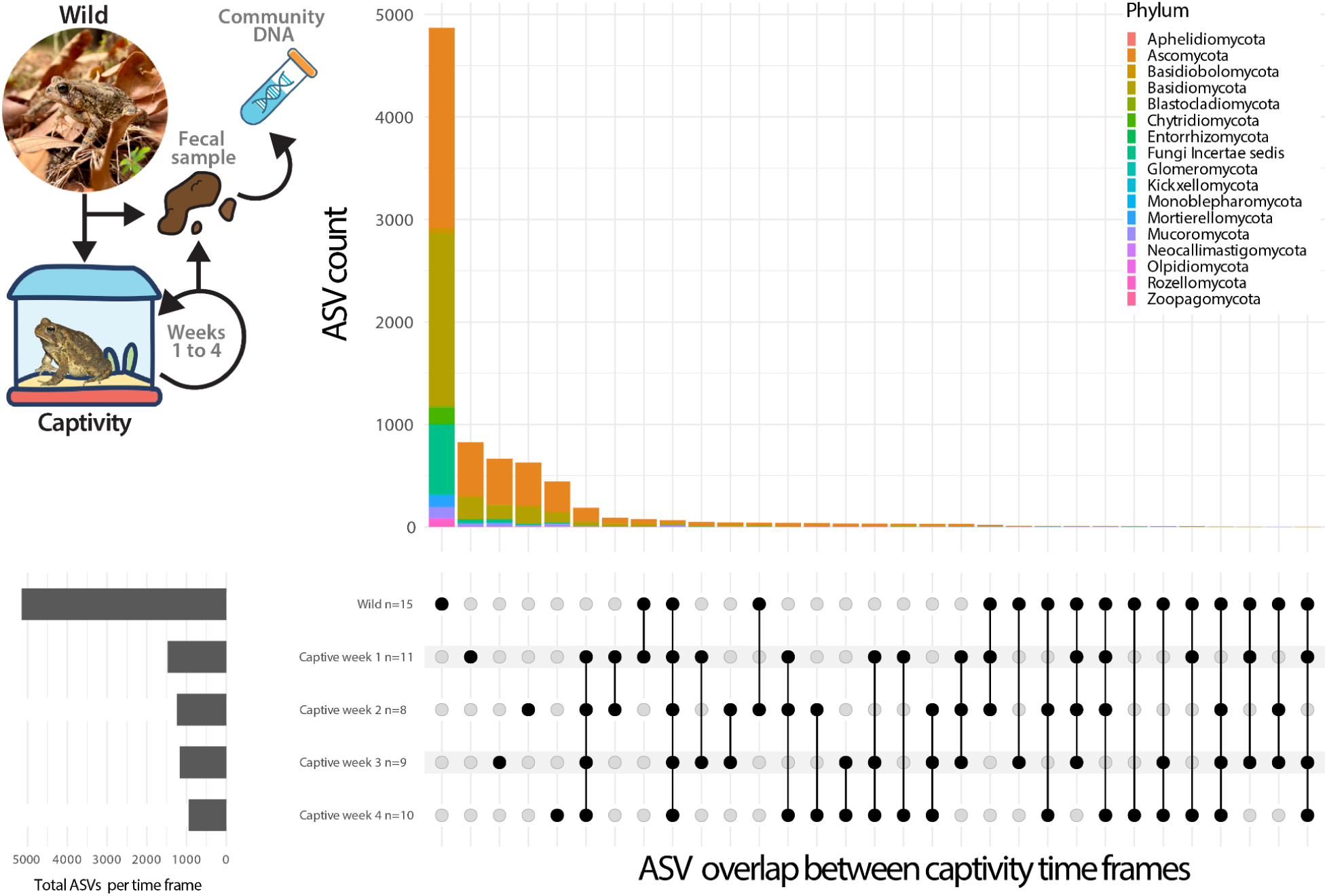
Assessing temporal dynamics of gastrointestinal fungal diversity pre- and post-captivity in Fowler’s Toad. Top: (Left) Depiction of the capture of toads from wild terrestrial environments, fecal sampling, and subsequent time in captivity. (Right) Stacked bar plot, with the Y-axis representing Amplicon sequencing variant (ASV) count and the X-axis indicating ASV count. Colors represent fungal phyla. **Bottom:** Left: Totals ASV count per sampling time frame. Right: Presence–absence dot plot where black dots mark ASV detection in each sample week and connecting lines trace the longitudinal occurrence of individual ASVs throughout the captive time series.

#### Metamplicon Sequencing and Broad Taxonomic Assessment

The ITS1 subunit was amplified by PCR and sequenced using 250X2 PE Illumina sequencing for 53 fecal pellets (wild n= 15, time series; wk1:n=11, wk2:n=8, wk3:n=9, wk4:n=10). Each sample was then processed using DADA2 (Callahan *et al*., 2016) and assigned taxonomic identifications based on the most recent UNITE fungal reference database (Abarenkov *et al*., 2024). A species accumulation curve analysis was conducted to determine if sufficient sampling had occurred to account for the recovery of Amplicon sequencing variants (ASV)s diversity (supplementary figure 2). ASVs represent a unique DNA sequence derived from amplicon sequencing data and can differ by single nucleotide changes, representing extremely fine measurements of taxonomic diversity. Species-accumulation curves for each treatment rose rapidly in wild populations, but diverged and flattened markedly between wild and captive groups (Supplementary Figure 2). All four captive timepoints (Weeks 1–4) reached an asymptote by 8–12 samples, with their 95 % confidence envelopes overlapping and yielding few additional ASVs beyond that threshold. In contrast, the wild-toad curve continued to rise even at 15 samples and showed no apparent plateau; its 95% envelope remained entirely above those of the captive groups (Supplementary Figure 2).

Across all samples, we identified 17,771 ASVs, representing 17 phylum-level assignments (Figure 1 and Supplementary Data). Despite these high numbers, 9,415 ASVs were assigned only to the Kingdom Level Fungi (53%), indicating the sheer undescribed diversity of the amphibian gut community. At the level of phylum, we recovered 8,356 ASVs, with the highest assigned group across all of our samples being the *Ascomycota* (n=4,214, 50%), followed by the *Basidiomycota* (n=2,508, 30%), *Mucoromycota* (n=242, 3%), and all other phyla representing the remaining ASVs (17% )(Figure 1). All deeper classifications are further reported in an interactive table provided in the Supplementary Data.

#### Characterization of alpha and beta diversity across captivity

Metrics of alpha diversity indicate an initial, steep decline in fungal diversity during the first week in captivity (Figure 2A and 2B). The diversity of the fungal microbiome appears to increase in the following weeks until week 4. We find significant differences when comparing the mycobiome of wild samples with those from captivity (PERMANOVA, F = 3.681, p < 0.001, Figure 2A). We see that the four-week time frame has the most distinct mycobiome compared to the wild. However, there are no differences in the mycobiome according to the sex of the sampled amphibians (F = 1.33, p > 0.5, Supp. Fig. 2).

**Figure 2.**
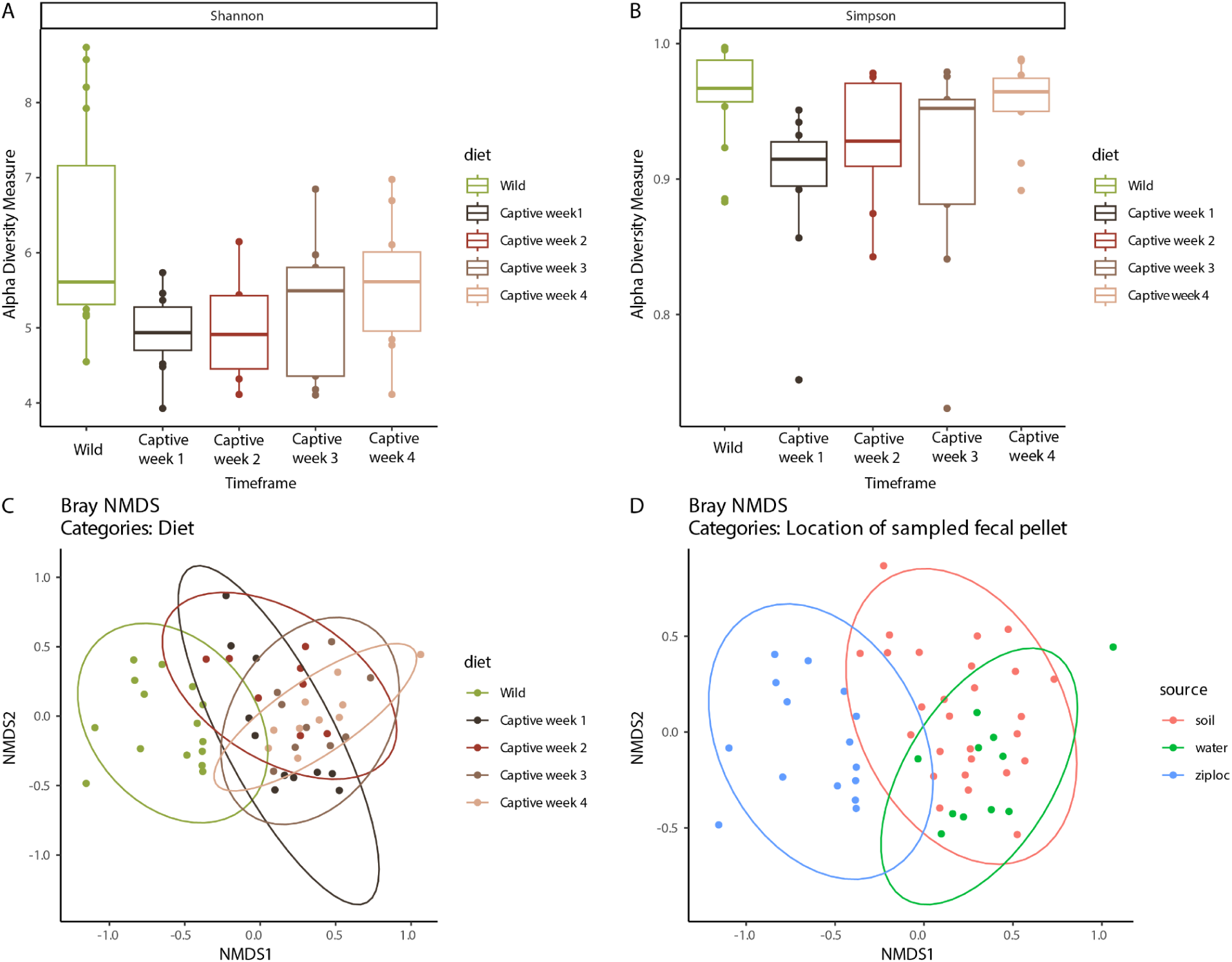
Diversity of Fowler’s Toad gut mycobiomes under wild and captive conditions. A) Shannon diversity index: Boxes show median and interquartile range, whiskers extend to 1.5× IQR, and points are individual samples. B) Simpson’s diversity index: Boxplots for the same groups and formatting as in (A). C). Bray–Curtis NMDS by time frame: Two-dimensional ordination of all wild and captive samples, colored by time point (same palette as A). Shaded ellipses are 95% confidence intervals for each group (stress = 0.14; PERMANOVA p < 0.01). D). Bray–Curtis NMDS by the physical location from which the pellet was collected in captivity. Ordination of captive samples only, colored by fecal-collection substrate, with 95% CI ellipses illustrating how environmental source influences community composition (stress = 0.12).

#### Temporal shifts in the mycobiome community during “wild” to “captive” transition, and the core mycobiome

A total of 453 ASVs differ significantly between the transition from wild to captive conditions (week 1). 102 ASVs show a significant increase in abundance in captivity, while 343 ASVs significantly decrease in abundance in captivity. Out of these, the phyla that increased the most in captivity were the *Ascomycota* (78 ASVs), followed by the *Basidomycota* (23 ASVs). There is more diversity in the phyla that decrease in captivity, where eight taxonomic categories are represented: *Ascomycota* (189 ASVs), *Basidomycota* (90 ASVs), *Basidiobolales* and *Mortierellomycota* (18 ASVs each), *Mucoromycota* (17 ASVs), and *Blastocladiomycota* with three ASVs.

Finally, 303 ASVs across seven phyla and 83 families represent the core mycogenome (e.g., species present across all stages of wilderness and captivity, Figure 3). Ascomycetes are the most represented with 138 ASVs, followed by *Basidiomycota* (47 ASVs), *Mucoromycota* (14 ASVs), *Basidiobolomycota* (4 ASVs), *Zoopagomycota* (3 ASVs), three ASVs recovered as *Incertae sedis*, and a single *Morteriellomycota* ASV.

**Figure 3.**
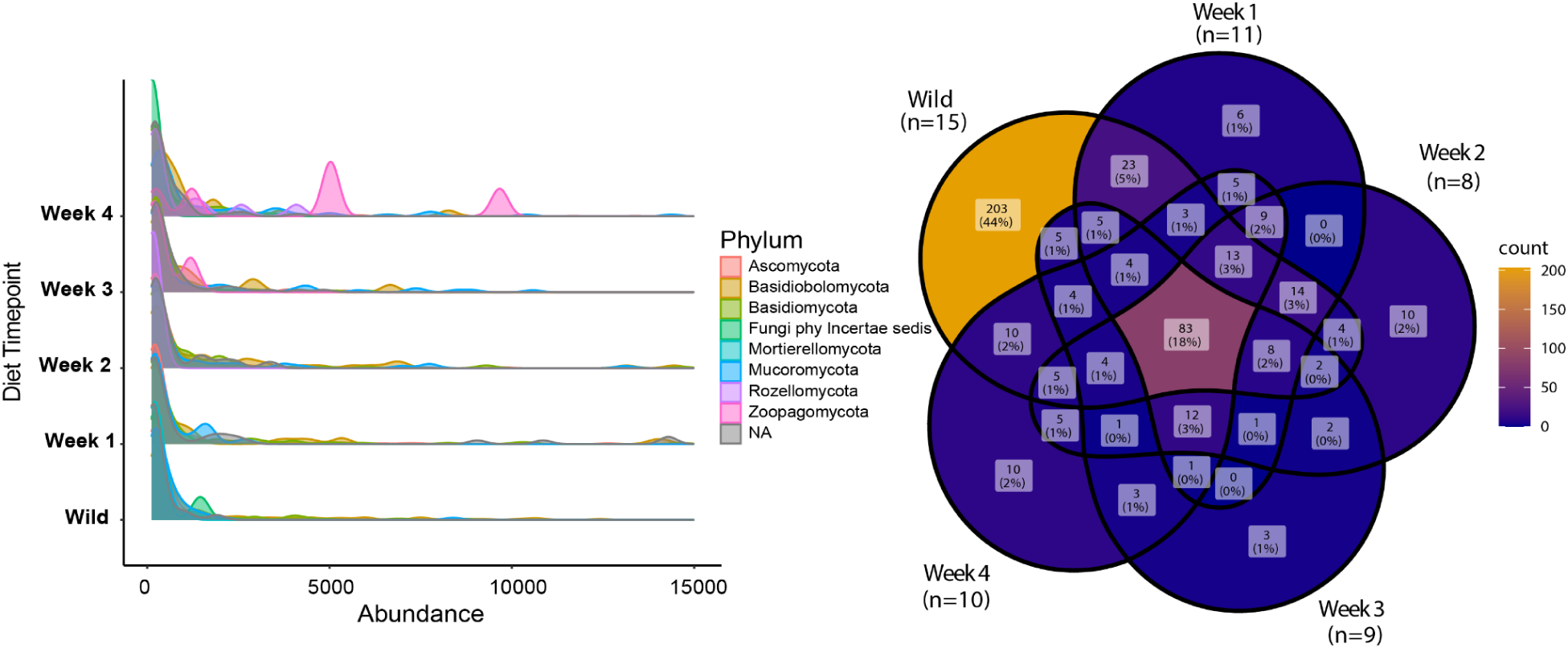
Phylum-level abundance distributions and Family-level ASV overlap across wild and captive timepoints. (Left) Ridgeline plots of Amplicon sequencing variant abundance by fungal phylum for wild toads (bottom) and weekly captive timepoints (Weeks 1–4, stacked upward). The x-axis shows raw ASV read counts, the y-axis lists diet conditions, and colors denote phyla. Peaks shifting in height and position illustrate how captivity reshapes the abundance landscape of each phylum. (Right) Five-set Venn diagram of ASV sharing among wild (n = 15) and captive toads at Weeks 1 (n = 11), 2 (n = 8), 3 (n = 9), and 4 (n = 10). Each circle represents a single time point; the numbers within each overlapping region indicate the count of ASVs and their percentage of the total ASV pool. This panel highlights the persistent “core” ASVs shared across all conditions and the transient taxa unique to specific weeks.

We observed significant differences in the decline of fungal phyla (Figure 3). In wild conditions, the most abundant fungal phyla were ascomycetes (41%), *Basidiobolales* (37%), *Basidiomycota* (14%), and *Mucoromycota* (4%). After one week in captivity, the mycobiome changed for some taxa, with a slight reduction for ascomycetes to 37%, a strong decrease for Basidiobolales to 19%, an increase of Basidiomycota to 35%, and Chytrids from 1% to 4%. *Mucoromycota* didn’t change. For week 4, Ascomycetes increased to 57% compared to the wild baseline, while *Mucoromycota* increased to 24%, and *Basidiomycota* and *Basidiobolales* decreased to 9% and 5%, respectively.

#### Association of Fungal Trophic Guild

There are marked differences in ecological (organisms grouped by how they acquire environmental resources) and functional guilds (organisms grouped by the general function they serve in an environment) assigned to the differentially abundant fungi (Figure 4). For the differentially abundant fungi that increase in captivity conditions, we find that Saprothophs (deriving nutrients from decaying biomatter) are found in the highest proportion, represented by 63 ASVs, followed by Patrotrophs (deriving nutrients from a pathogenic relationship with a host )/Saprothrophs (15 ASVs), Patrotrophs/Saprothrophs/symbiotrophs (deriving nutrients from a symbiotic relationship with a host) (11 ASVs), and Patrotrophs (4 ASVs). Once again, there is a larger diversity in the differentially abundant ASVs in the wild samples, where nine guild categories are represented: Saprothophs with 91 ASVs, followed by Patrotrophs/Saprothrophs (81 ASVs), Patrotrophs/Saprothrophs/symbiotrophs (64 ASVs), Saprotroph-Symbiotroph (30 ASVs), Pathotroph (17 ASVs), Pathotrophs (3 ASVs), Pathotroph-Symbiotrophs (3 ASVs), and Symbiotrophs (2 ASVs). No Symbiotrophs were represented in the differentially abundant ASVs from captivity.

**Figure 4.**
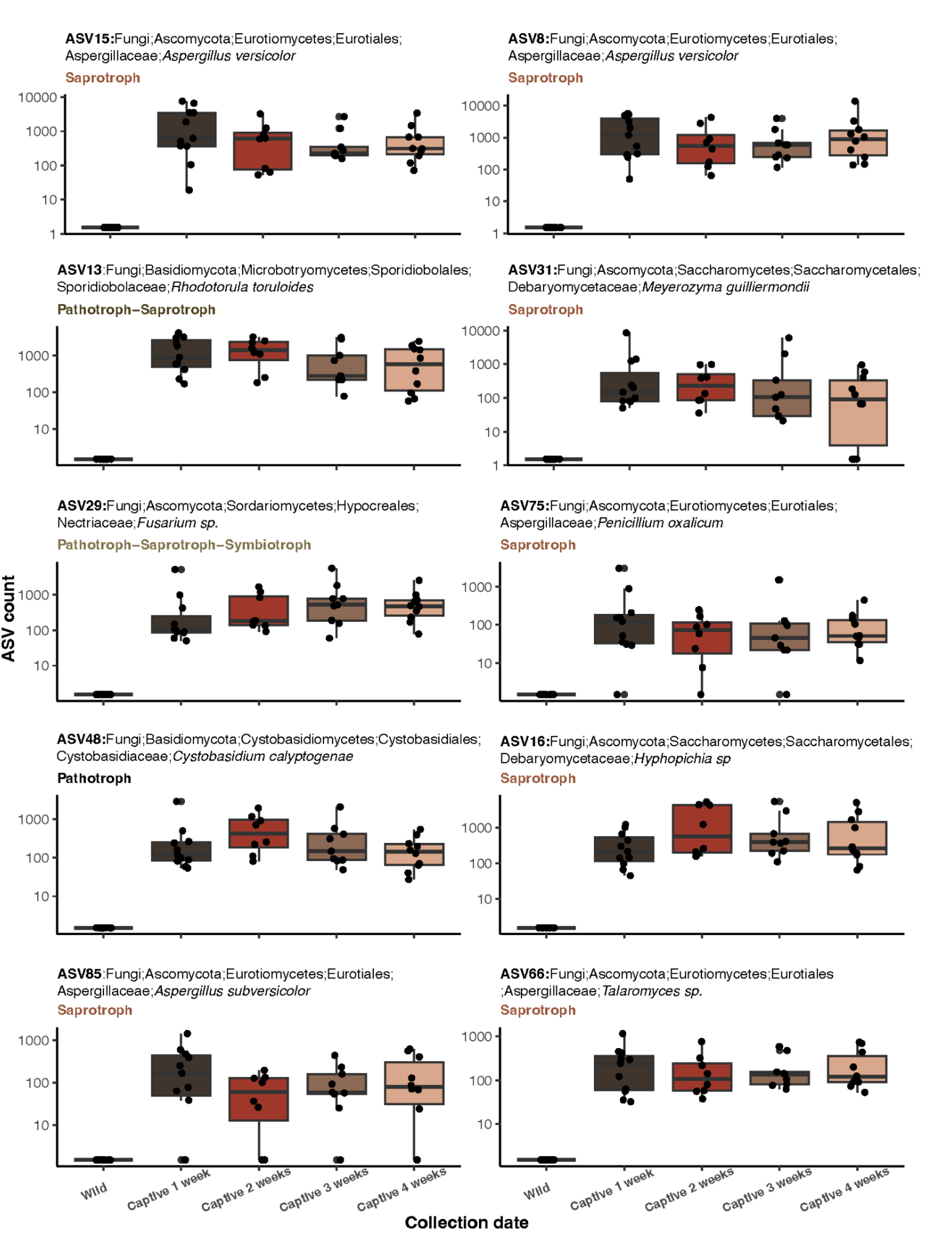
Captivity-enriched fungal ASV dynamics. Log₁₀ ASV count for a subset of the ten ASVs with the largest mean increase before and after captivity. Each ASV is faceted by FUNGuild ecological guild (saprotroph, pathotroph–saprotroph, pathotroph–saprotroph–symbiotroph, pathotroph). Points represent individual samples (n = 9–15).

## Discussion

Our study examined the static core microbiome and the dynamic community shifts that occur as amphibians transition from their natural to captive environments. Using the common Fowler’s toad, we aimed to understand how captivity impacts fungal diversity, taxonomic composition, and functional guilds associated with amphibian mycobiomes. Our results reveal a stable “core” mycobiome and dramatic, captivity-driven shifts in the GI-associated Fungi of Fowler’s Toads, emphasizing the need to account for these dynamics in amphibian conservation. We detected a small suite of fungi that persisted across wild and captive conditions. In contrast, the broader mycobiome underwent dramatic restructuring: fungal richness plummeted by over 50% within one week of captivity, followed by a partial rebound over the next month. This temporal pattern suggests an acute microbiome disruption upon transfer to captivity, with some opportunistic taxa recovering as the host and environment stabilize.

The early plateau of the captive accumulation curves confirms that our sampling depth was sufficient to capture the depleted mycobiome present under captive conditions. In contrast, the unabated rise of the wild curve reveals a substantially richer fungal community in free-living toads (Supp. Fig 2). These results suggest that this disparity is a genuine biological effect, reinforcing the conclusion that captivity imposes significant selective pressures on host-associated fungi, thereby dramatically reducing amphibian gut mycobiome diversity. We acknowledge that the accumulation curves (Supplementary Figure 2) for the wild population show under-sampling, which is not ideal for characterizing fungal community richness and composition. However, these limitations were expected, as they are the results of an initial trial and comparison of the diversity losses between wilderness and captivity. This comparative framework is therefore critical for highlighting the pronounced loss of microbial diversity in *ex situ* conservation and guiding future efforts to monitor and mitigate mycobiome erosion in at-risk amphibian populations with larger sample sizes in future studies.

### The “core” mycobiome and its potential role

The core mycobiome, defined here as the 308 ASVs found in every wild and captive sample, was dominated by Ascomycetes (134 ASVs), with substantial contributions from *Basidiomycota* (36 ASVs), *Mucoromycota* (14 ASVs), *Chytridiomycota* (6 ASVs), *Basidiobolales* (4 ASVs), and *Mortierellomycota* (1 ASV). At the genus level, *Basidiobolus* (20%), *Mucor* (15%), *Aspergillus* (7%), and *Papiliotrema* (5%) comprised the majority of this persistent community. These genera are known for their presence in the GI tract of vertebrates. For example, *Basidiobolus* species form symbiotic relationships with amphibians and have been consistently found across different geographic locations and hosts (Walker *et al*., 2020; Vargas-Gastélum *et al*., 2024; Hincher *et al*., 2025) , although the role they play as symbionts remains unknown. Likewise, core Ascomycetes, such as *Aspergillus,* are renowned for their carbohydrate-degrading enzymes (De Vries & Visser, 2001), which may complement host digestion. Meanwhile, *Mucor’s* rapid growth under fluctuating moisture hints at its resilience to substrate shifts in captivity. Together, this conserved assemblage could underpin essential digestive, immunological, and perhaps even behavioral functions, providing a microbial “backbone” that persists despite the perturbations of artificial environments, which has been shown in other model organisms such as mice (Hanski *et al*., 2024) and endangered species, such as the Giant Panda (Huang *et al*., 2023). By characterizing these steadfast fungal partners, we establish a baseline against which the health and stability of amphibian mycobiomes can be assessed in both conservation and husbandry contexts.

### Temporal shifts of the mycobiome after captivity

The overall fungal community composition experienced marked shifts upon transition to captivity, particularly in the dominant fungal phyla. Temporal results, in more detail, showed a significant reduction in fungal diversity within the first week of captivity, followed by a partial recovery over the subsequent four weeks. Ascomycetes remained the most prevalent fungal group across all conditions, though their relative abundance increased over time in captivity. Basidiobolales showed a steep decline from 37% in wild conditions to just 5% by the fourth week of captivity. This suggests that Basidiobolales may be sensitive to environmental changes associated with captivity, potentially due to altered host physiology, diet, or habitat conditions. Basidiomycota exhibited a transient increase in the first week of captivity but declined significantly by the fourth week (Figure 3A). This initial increase could indicate opportunistic colonization in response to the novel captive environment, followed by competitive exclusion by other taxa. *Mucoromycota*, which contains microfungi with broad ecological guilds, including saprophytic and pathotrophic, showed a steady increase over time. The noticeable shift of ASVs assigned to *Mucoromycota* with FUNguild assessments of Saprotroph (n=231/242) versus Saprotroph-Symbiotroph (n=11/242) suggests that captivity may provide conditions favorable for this group, possibly through shifts in moisture levels or substrate availability (Figure 3A), or through an increase in niche space due to the overall decrease of the mycobiome community. *Chytridiomycota*, though initially present at low abundance, increased slightly in captivity, indicating potential responses to environmental or host-associated factors (Figure 3A). These taxonomic shifts suggest that captivity imposes intense selective pressures on the amphibian mycobiome, leading to the loss of specific fungal taxa while favoring others, particularly Ascomycetes and Mucoromycota.

These microbial shifts have been well-documented in captive populations of multiple host taxa, including birds such as the Brown Kiwi (San Juan *et al*., 2021), primates (Wang *et al*., 2021), and large mammals like the black rhinoceros (Gibson *et al*., 2019). Despite this growing body of research, significant gaps remain in our understanding of how captivity influences microbial communities in less-studied taxa, such as amphibians. Additionally, while bacterial microbiomes have been extensively investigated, the mycobiome has largely been overlooked, leaving its diversity, ecological functions, and host interactions poorly understood. Fowler’s Toads are abundant and widespread (IUCN, 2023), making them a suitable model for examining conservation strategies that can then be applied to at-risk species (Poo & Hinkson, 2020; Poo *et al*., 2022; Malter *et al*., 2025)

### Variation of fungal ecological guilds after captivity

Assigning ecological guilds with FUNGuild (Nguyen *et al*., 2016) revealed a stark functional restructuring of the toad mycobiome under captivity. Among ASVs significantly enriched in captivity, saprotrophs dominated with 63 ASVs, followed by pathotroph–saprotrophs (15 ASVs), pathotroph–saprotroph– symbiotrophs (11 ASVs), and pure pathotrophs (4 ASVs). In contrast, ASVs that declined from the wild encompassed nine guilds, led by saprotrophs (91 ASVs), pathotroph–saprotrophs (81 ASVs), and pathotroph–saprotroph–symbiotrophs (64 ASVs), but also including saprotroph–symbiotrophs (30 ASVs), pure pathotrophs (17 ASVs), and even rare pure symbiotrophs (2 ASVs). Notably, no ASVs classified solely as symbiotrophs were differentially abundant between nature and captivity, indicating there is little difference in the abundance of dedicated mutualists under both natural *ex situ* conditions, but a strong pattern of symbiotrophs that have guild plasticity. The shift toward saprotrophic and pathotrophic lifestyles, and away from symbiotic modes, suggests that dietary simplification and altered substrates in captivity favor generalist decomposers, potentially at the expense of fungi that provide host-beneficial services such as nutrient provisioning or pathogen defense. Maintaining functional guild diversity, therefore, may be as critical as preserving taxonomic diversity in captive husbandry and reintroduction programs.

### The importance of understanding the shifts in mycobiome composition as part of restoration and conservation efforts

Building on our trial data, we can leverage the temporal patterns of diversity loss and taxonomic turnover we observed in Fowler’s Toads to investigate “microbial crashes” in other at-risk amphibians. For example, eastern Hellbender salamanders (*Cryptobranchus alleganiensis*), one of North America’s largest amphibians, face significant population declines across much of their range due to habitat loss, water pollution, disease, and stream sedimentation. Although extant in distinct populations, they are considered near-threatened or endangered in many states, and the eastern hellbender was recently listed as vulnerable due to decreasing population numbers, which are tending toward endangered status (IUCN, 2021). Conservation efforts are underway to help stabilize and restore populations, including habitat restoration, streambank stabilization, captive breeding, and head-starting programs, where juveniles are raised in captivity before release (Macklem *et al*., 2025). By sampling hellbenders at key stages of head-starting and captive-breeding programs, we can test whether similar rapid losses of fungal diversity occur upon transfer to artificial stream enclosures and whether particular taxa (e.g., core *Ascomycetes* or *Basidiobolales*) resist collapse or predict reintroduction success.

Similarly, our framework can be applied to critically-endangered species that rely heavily on captive-release programs, such as the Wyoming Toad (*Anaxyrus baxteri*) and dusky gopher frogs (*Lithobates sevosus*). In these cases, captive-bred individuals that are naive to the complexity of their natural habitats are released into the field to build new wild populations (Odum RA, 2005; Bogisich *et al*., 2025). Monitoring shifts in mycobiome structure across life stages and release sites could pinpoint fungal partners essential for larval and juvenile development and pathogen defense, informing husbandry adjustments (e.g., substrate choices or probiotic applications) that preserve functional guilds. Ultimately, transferring our insights from Fowler’s Toads to these conservation and species recovery programs will help identify microbial indicators of host health, optimize *ex situ* rearing conditions, and improve survival rates upon release, which contribute toward mitigating amphibian declines worldwide.

## Conclusion

Our study investigates the dynamic nature of the amphibian mycobiome and the profound selective pressures that captivity imposes on fungal diversity, community structure, and functional guilds. The rapid declines and compositional shifts we observed reveal how *ex situ* conditions can destabilize host-associated microbial assemblages, potentially impacting amphibian health, immunity, and reintroduction success. Future work should assess the long-term physiological and fitness consequences of captivity-driven mycobiome changes, particularly within conservation, captive-release, and head-start programs, to fully understand these impacts. More broadly, unraveling the intricate relationships between fungi and their amphibian hosts under varying environmental contexts will be essential for designing husbandry and management practices that preserve microbial stability. Ultimately, integrating mycobiome monitoring and manipulation into *ex situ* conservation strategies may prove critical for bolstering the resilience and survival of both captive and wild amphibian populations.

## Methods

### Animal capture and care

Animal handling and care procedures were approved by the Memphis Zoo Institutional Animal Care and Use Committee (Approval 2022-04) and the Tennessee Wildlife Resource Agency (Permit 2430), which are described briefly below. In May 2022, fifteen adult Fowler’s Toads were captured by hand in relatively even sex ratios (Female=7, Male =6, Unknown=2) in Shelby County, Tennessee (USA). Toads were located between 2100 and 2400 hours through visual encounter surveys using a headlamp with a brightness of 1,250 lumens. The sex of captured toads was determined by the presence of a darker throat in males, a secondary sexual characteristic present in males during the breeding season. Toads were housed in same-sex groups of four individuals in 10-gallon glass aquariums (50.8 cm L × 25.4 cm W × 30.48 cm H) outfitted with coconut shavings, tap water that was dechlorinated and conditioned using API Tap Water Conditioner, and a hide box for cover. Daily water changes were conducted and spot cleaning of excrement was performed, with substrate replacement occurring as needed. The provided diet consisted of a combination of crickets, mealworms, and superworms offered four times per week. Artificial light was provided in a 12:12 h light-to-dark cycle, and the room was maintained at ∼23 ^0^C. All materials and tools used during the study were cleaned with a diluted 15% bleach solution and thoroughly rinsed with tap water before use.

### Fecal collections, DNA Extraction, quantification, and control

Fecal samples were collected from toads by scooping the fecal sample into a 2 mL Eppendorf tube and stored frozen at -20 °C until ready for processing. Total gDNA was extracted using the QiAmp PowerSoil Pro DNA Kit (Qiagen, Cat. No. 51804) using the manufacturer’s suggested protocol with the specific modifications of initial vortexing time of the fecal sample increased to 15 minutes to ensure homogeneity with the C1 (lysis) buffer, and the C3 (wash) and C6 (elution) buffers were incubated at 65°C before use in a water bath.

DNA concentrations for each sample were measured with a Qubit 4 Fluorometer using the 1x dsDNA High Sensitivity reagents (ThermoFisher Scientific). Sample purity was also checked at 260, 280, and 230 nm with a Nanodrop One Microvolume UV-Vis Spectrophotometer (ThermoFisher Scientific). All gDNA extracts were stored at -20°C until they were ready for metabarcode sequencing. Fecal pellets were collected weekly from both male and female toads, resulting in a total of 53 samples for sequencing.

### Metabarcode sequencing and taxonomic classification

To characterize the microbial community, we generated metamplicon data of the ITS1 subregion of the fungal internal transcribed spacer region, considered one of the most reliable housekeeping genes for fungal species identification (Schoch *et al*., 2012). The first round of amplification was performed for each sample using the ITS5-1737F (GGAAGTAAAAGTCGTAACAAGG) and ITS2-2043R primers (GCTGCGTTCTTCATCGATGC). Amplified sequences were then sent to NovoGene Inc. (Sacramento, CA), which performed the secondary amplification and indexing. Pooled samples were then sequenced using PE 2X250 Illumina libraries on a Novaseq. Raw sequencing statistics are reported in Table 1. Demultiplexing was also performed at Novogene using their in-house bioinformatic service. Raw reads were processed in R using sequence trimming specific to the variable ITS region using cutadapt v3.4 (Martin, 2011), and the DADA2 pipeline to produce amplicon sequence variants (ASVs) with standard settings (Callahan *et al*., 2016). Taxonomic classification of ASVs was performed with the UNITE database (sh_general_release_dynamic_04.04.2024) (Nilsson *et al*., 2019; Abarenkov *et al*., 2024).

**Table 1.**
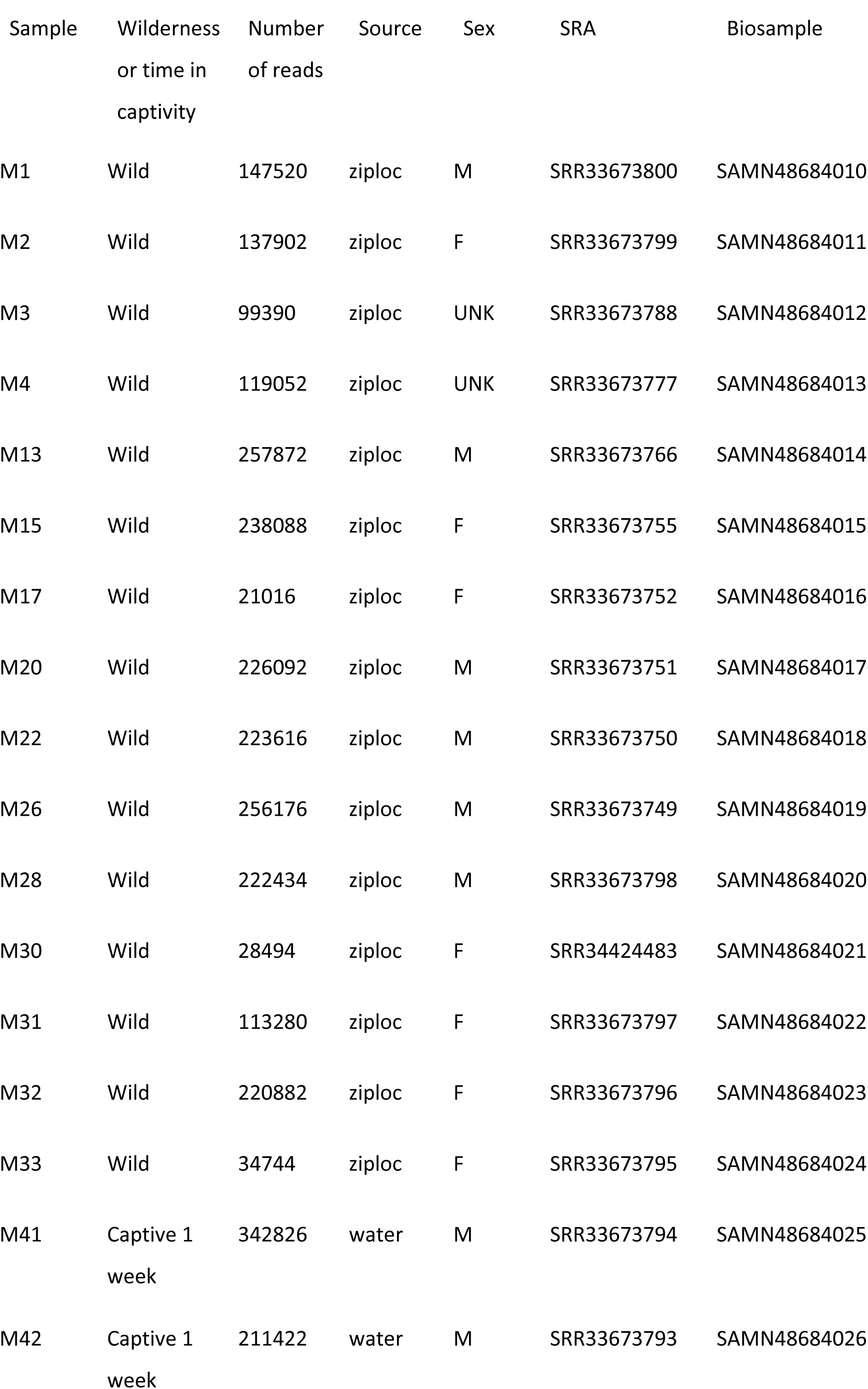
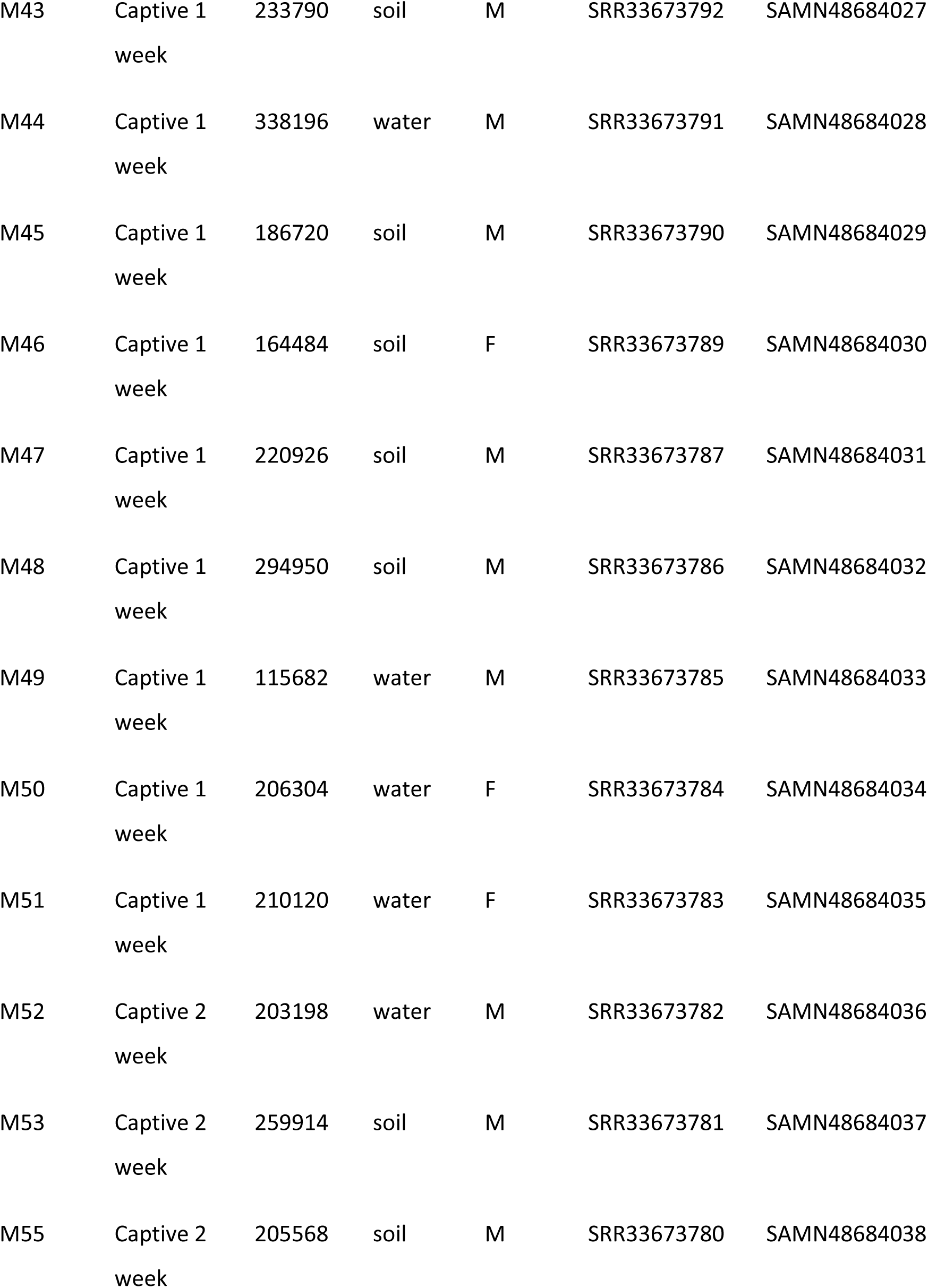
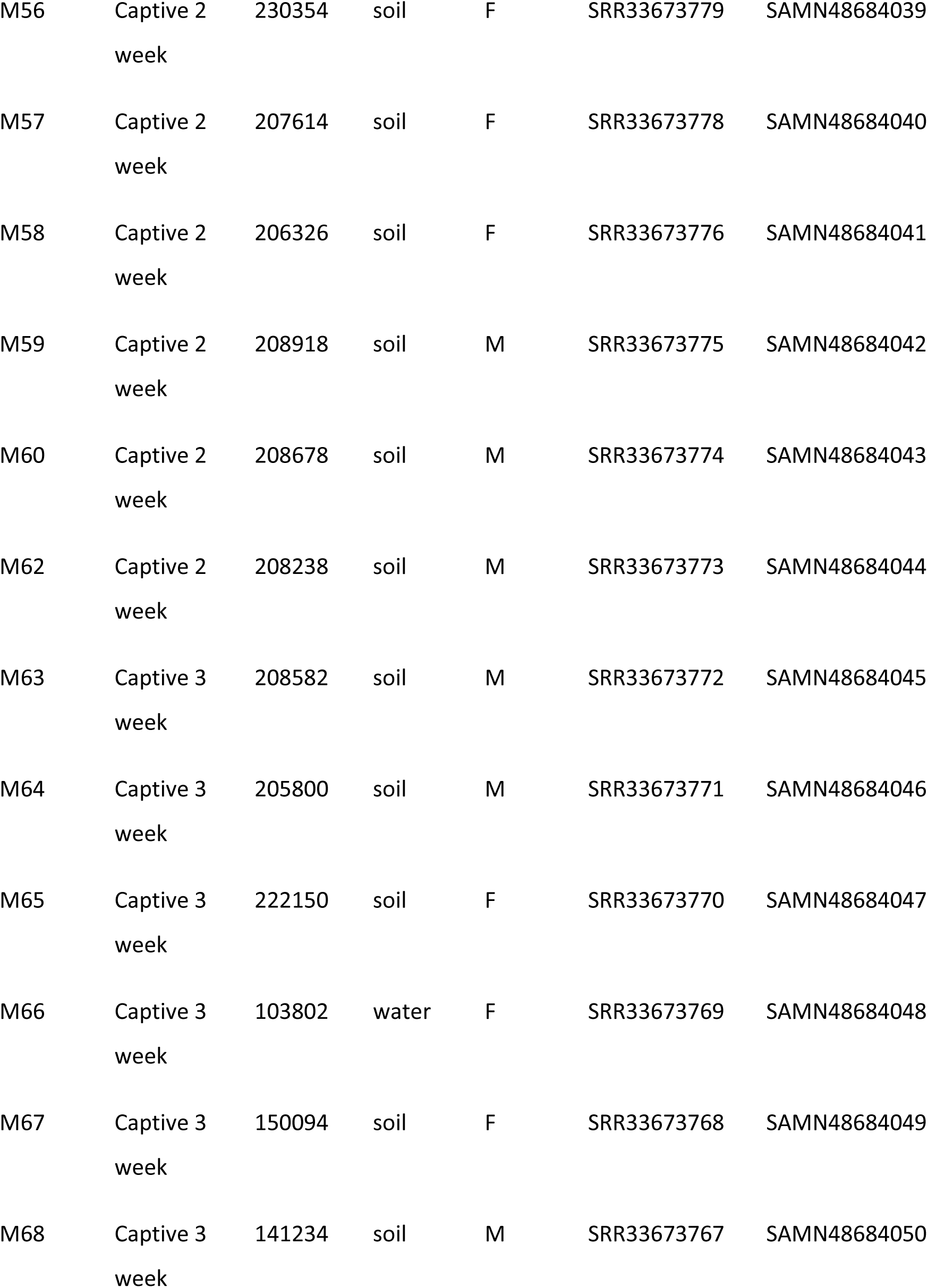
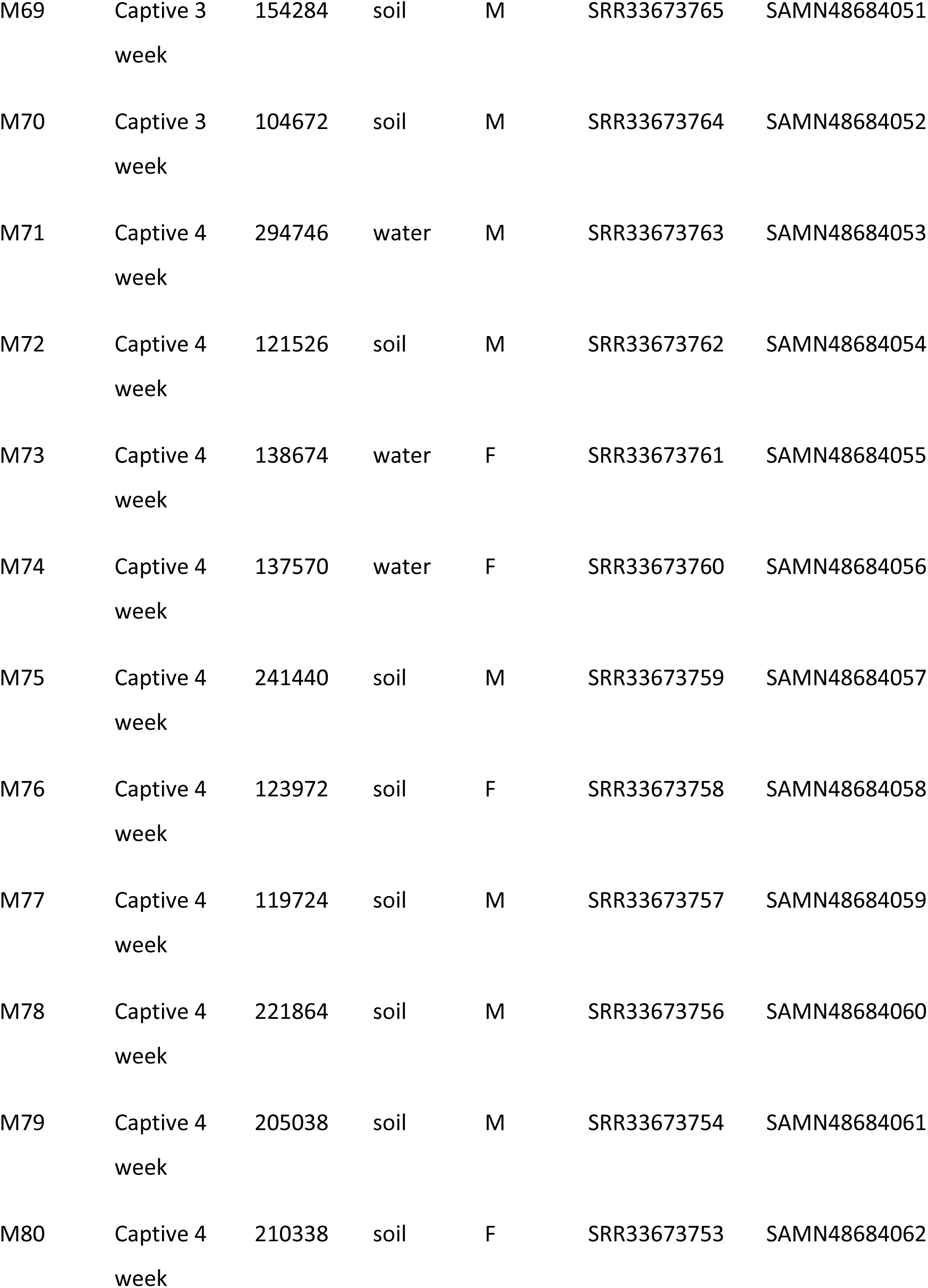
Sample information and sequencing statistics.

### Data exploration and visualization

Data exploration, visualization, and generation of interactive ASV tables (supplementary data) were performed using the R package MiscMetabar v0.14.2 (Taudière, 2023). Preprocessing of our phyloseq object was first performed using the function clean_pq with simplify_taxo=TRUE to clean and simplify the taxonomic classifications within our OTU matrix. An overall visual summary for our phyloseq object was created using the summary_plot_pq function (Supplementary Figure 1). To generate the UpSet plot (Figure 1) to investigate phyla overlap between diet time frames, we first subset our phyleq object using the ps_clean function from MiscMetabar at the Phylum level, removing all NA designations. The upset_pq of MiscMetabar was then used with a minimum number of sequences set to 2 to remove single ASVs. Ridge plot analysis (Figure 3, Left) was performed using MiscMetabar’s ridges_pq function with an alpha value of 0.5 to visualize spikes in specific phyla based on diet time frames. To visualize the overlap of Family-level ASVs across all diet time frames, a Venn diagram (Figure 3, Right) was generated with the ggvenn_pq function of MiscMetabar with the options taxonomic_rank “Family”.

### Species accumulation analysis

A species accumulation plot was generated from our phyloseq object using the specaccum function of the R package VEGAN v2.6-10 using a random sampling method with 100 permutations (Dixon, 2003). Samples were grouped by diet time frames, and 95% ASV recovery thresholds were quantified and plotted using ggplot2 v3.5.2 (Wickham, 2016) (Supplementary Figure 2).

### Statistical and biodiversity measurements and fungal trophic guild determination

Community analyses were performed using the R package Phyloseq V1.44.0 (McMurdie & Holmes, 2013). Visualizations, including graphs and figures, were plotted using the data visualization package ggplot2 (Wickham, 2016). The Shannon diversity index, used to assess alpha diversity, and the Bray-Curtis dissimilarity index, used to evaluate beta diversity, were employed to analyze differences in community composition. This approach considered relative abundances, ignored shared absences, and effectively captured ecological gradients, making it ideal for analyzing microbial community shifts across our treatments. The identification of differentially abundant taxa before and after captivity was determined using DESeq2 (Love *et al*., 2014). The variance-stabilizing transformation (VST) was used to normalize count data, and differentially abundant taxa were identified with the DESeq function (adjusted p-value < 0.01). Log2 fold changes were calculated for significant taxa enrichment or depletion in fecal samples collected before placement in captivity and at each subsequent week. A quality control cutoff was introduced to reduce noise in the dataset by only including taxa if a log-fold change in abundance greater than two and an adjusted p-value of 0.01 was observed. Taxa that could not be assigned (NA) at the phylum level were excluded from downstream analysis as they provided no distinct information to be evaluated. In addition to the taxonomic assignment, the ecological guild, which has been shown to provide a broad estimation of what a fungal community may be doing within an environment, was assigned to ASVs using the FUNGuild database v1.1 (Nguyen *et al*., 2016). Significant differences in trophic mode abundance were analyzed by comparing wild samples to all captivity weeks and vice versa, and plotted using ggplot2 (Wickham, 2016).

## Acknowledgements

We thank the Memphis Zoo for its commitment to wildlife conservation and Dr. Steve Reichling for his support in pursuing research questions that provide insights into the care and management of animals in captivity.

## Author Contributions

**Alexander J. Bradshaw** performed data analysis, figure creation, and manuscript preparation and editing of the manuscript. **Sinlan Poo** collected samples, maintained specimens in captivity, and prepared and edited the manuscript. **Anne Devan-Song** collected samples and prepared and edited the manuscript.

**Tracey E. Malter** collected samples, maintained specimens in captivity, and contributed to the manuscript. **Rylie M. Strasbaugh** collected samples, maintained specimens in captivity, and contributed to the manuscript. **Brianna Bodner** performed fecal DNA extraction and quality control. **Madison R. Hincher** performed fecal DNA extraction and quality control**. Javier F. Tabima** performed data analysis, figure creation, and preparation and editing of the manuscript.

## Data availability

All raw sequencing data have been deposited in the Short Read Archive (SRA) under Bioproject number PRJNA1266671, including Individual SRA and Biosamples (Table 1). All supplementary data, including the full Rmd file and phyloseq objects, as well as an interactive ASV table, can be downloaded from the Open Science Framework (OSF) under DOI: 10.17605/OSF.IO/5ANZ2 Any code or specific script requests should be sent to the corresponding author.

---

**Fig. S1:**
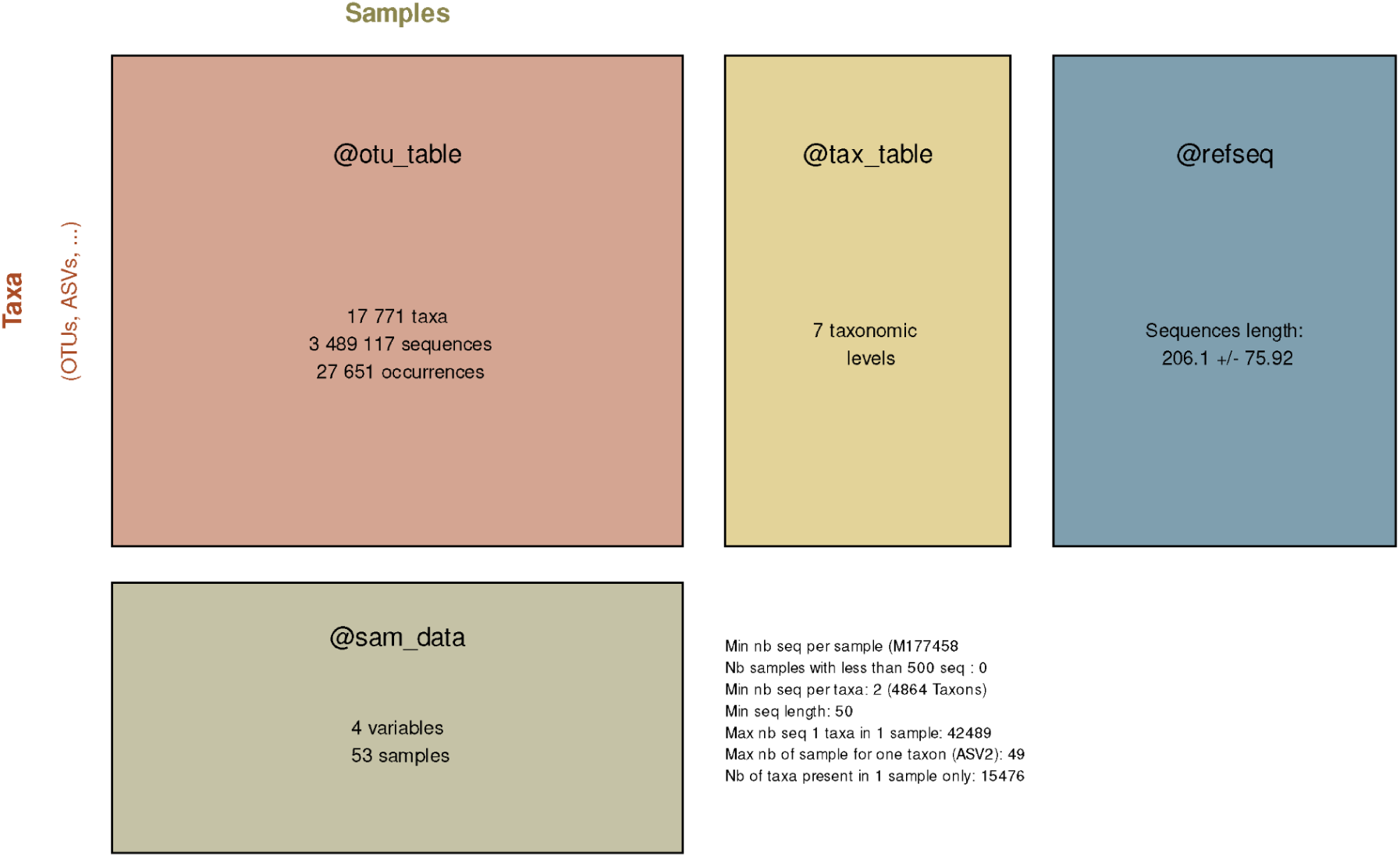
Visual summary of Phyloseq object statistics. Visual summary of the phyloseq object representing all raw data from this work. Separate data matrices are represented by color squares and labeled with their designation within the phyloseq object. Primary stats are reported for the total phyloseq object adjacent to @sam_data.

**Fig. S2:**
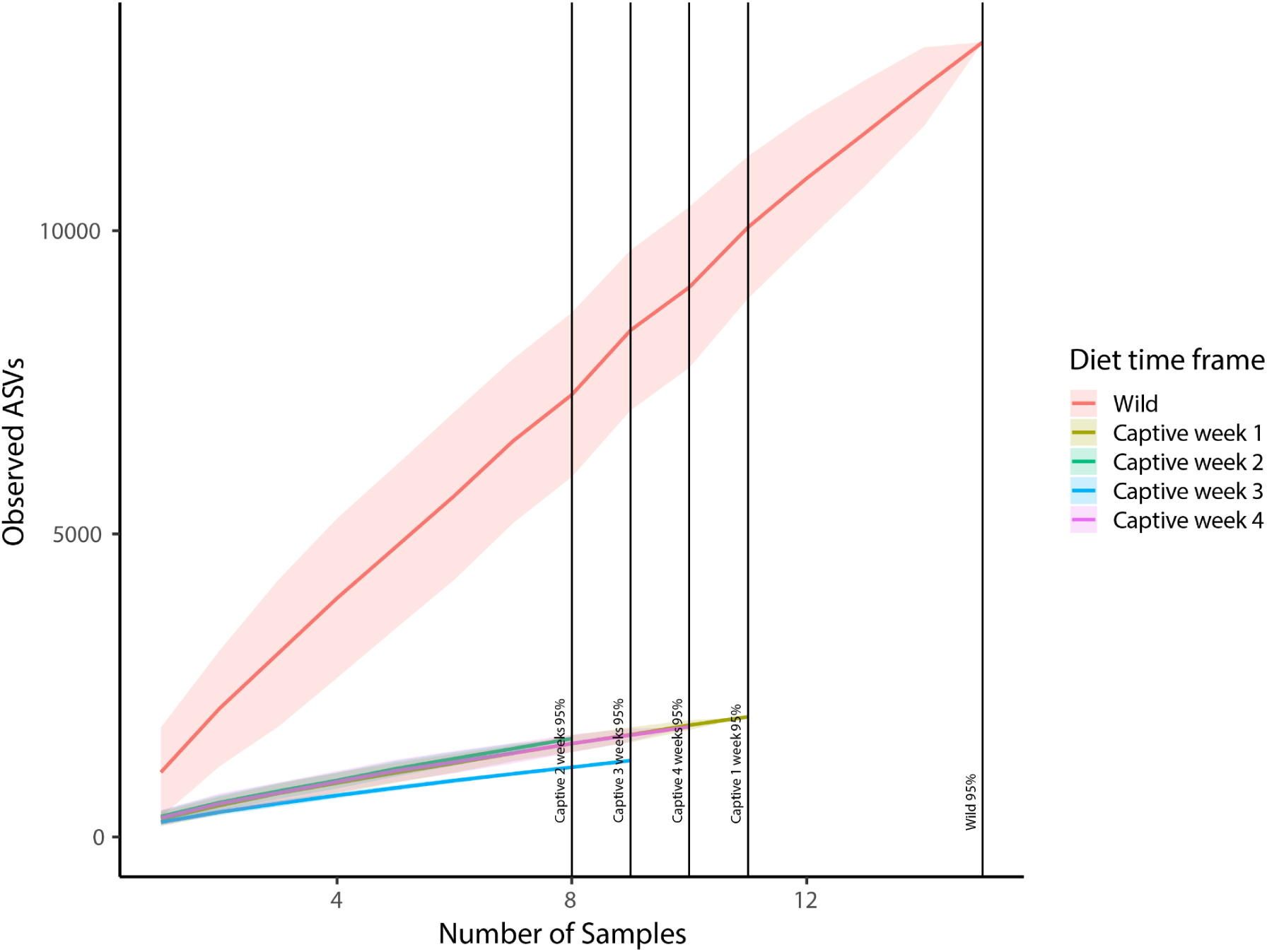
Species-accumulation curves of gut fungal ASV richness in Fowler’s Toads under wild and captive conditions. Mean cumulative ASV richness (y-axis) is plotted against the number of fecal samples (x-axis) for wild toads and individuals in captivity. Shaded bands represent 95 % confidence intervals around each curve. Vertical lines indicate the sample size at which each group reaches 95 % of its asymptotic richness. Wild toads display a continuously rising curve without a plateau, whereas all captive groups plateau by 8–12 samples.

**Fig. S3.**
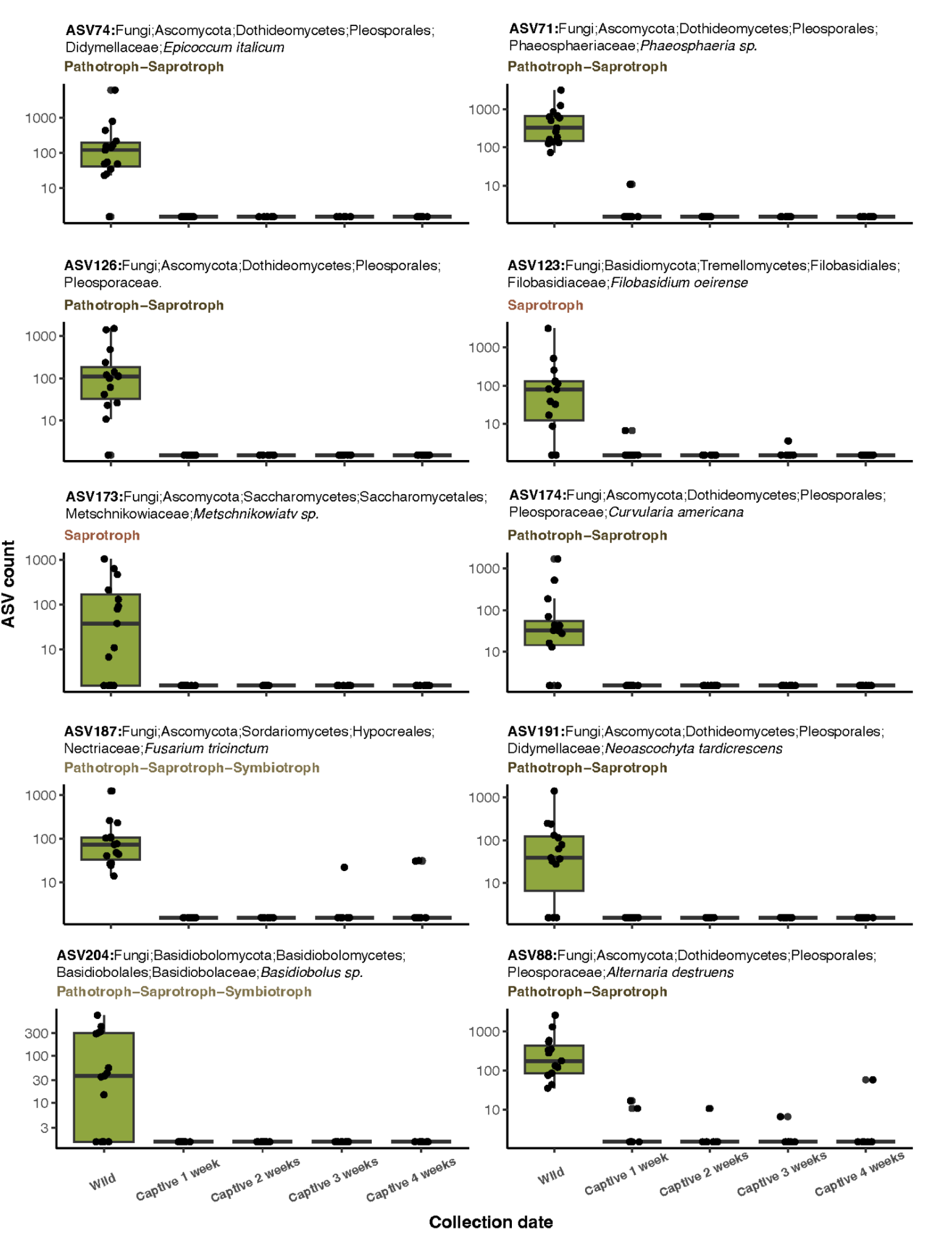
Captivity-enriched fungal ASV dynamics. Log₁₀ ASV count for a subset of the ten ASVs with the largest mean decrease under captivity faceted by FUNGuild ecological guild (saprotroph, pathotroph–saprotroph, pathotroph–saprotroph–symbiotroph, pathotroph). Points represent individual samples.

